# Cerebral White Matter Sex Dimorphism in Alcoholism: A Diffusion Tensor Imaging Study

**DOI:** 10.1101/229609

**Authors:** Kayle S. Sawyer, Nasim Maleki, George Papadimitriou, Nikos Makris, Marlene Oscar-Berman, Gordon J. Harris

## Abstract

**Background:** Excessive alcohol consumption is associated with widespread brain damage, including abnormalities in frontal and limbic brain regions. In a prior study of neuronal circuitry connecting the frontal lobes and limbic system structures in abstinent alcoholic men, we demonstrated decreases in white matter fractional anisotropy (FA) on diffusion tensor magnetic resonance imaging (dMRI). In the present study, we examined sex differences in alcoholism-related abnormalities of white matter connectivity.

**Methods:** dMRI scans were acquired from 49 abstinent alcoholic individuals (26 women) and 41 nonalcoholic controls (22 women). Tract-based spatial statistical tools were used to estimate regional FA of white matter tracts and to determine sex differences and their relation to measures of alcoholism history.

**Results:** Sex-related differences in white matter connectivity were observed in association with alcoholism: Compared to nonalcoholic men, alcoholic men had diminished FA in portions of the corpus callosum, the superior longitudinal fasciculi II and III, and the arcuate fasciculus and extreme capsule. In contrast, alcoholic women had higher FA in these regions. Sex differences also were observed for correlations between corpus callosum FA and length of sobriety.

**Conclusions:** Sexual dimorphism in white matter microstructure in abstinent alcoholics may implicate underlying differences in the neurobehavioral liabilities for developing alcohol abuse disorders, or for sequelae following abuse.

## Introduction

Excessive alcohol consumption is associated with widespread abnormalities in brain regions involved in cognition and emotion, as evidenced by multiple neurobehavioral, neuroimaging, and post-mortem studies (Oscar-Berman *et al*, 2014; Oscar-Berman and Marinković, 2007). However, the majority of neuroscientific studies in alcoholism have focused on men (Ruiz and Oscar-Berman, 2013; Susan Mosher Ruiz, 2015), and there is limited scientific literature evaluating sex-specific similarities and differences in brain pathology associated with alcoholism. Moreover, there is accumulating evidence of sex differences in pharmacokinetic responses to alcohol consumption and metabolism (Chrostek *et al*, 2003; Ramchandani *et al*, 2001), which may further impact sex differences in health problems mediated by social and environmental, as well as neurobiological, physiological, and genetic factors (Ceylan-Isik *et al*, 2010; Verhulst *et al*, 2015).

Multiple studies, focused mainly on alcoholic (ALC) men, have reported significant structural brain abnormalities that are correlated with indices of alcoholism history, such as the duration of alcoholism, the amount of alcohol consumed, or the duration of abstinence (Le Berre *et al*, 2015; Segal *et al*, 2016). These abnormalities involve cortical gray matter as well as white matter fiber tracts, and may appear before behavioral symptoms arise. Alcoholism is particularly damaging to cerebral white matter, as has been revealed by post-mortem neuropathology (Harper *et al*, 2003; Putzke *et al*, 1998; Tarnowska-Dziduszko *et al*, 1995) and by *in vivo* structural MRI studies (Pfefferbaum *et al*, 1992, 2009). Consistent with these findings, post-mortem RNA analyses of superior frontal lobe samples found that genes related to myelin structure were down-regulated in alcoholics (Lewohl *et al*, 2000).

In our prior work, we used diffusion tensor magnetic resonance imaging (dMRI) to examine the integrity of white matter fiber tracts in the brains of abstinent ALC men compared with nonalcoholic control (NC) men (Harris *et al*, 2008). We found that the ALC men had lower fractional anisotropy (FA) than NC men within several white matter fiber tracts. In another study we found that ALC men had smaller volumes of brain regions in the extended reward and oversight system (Makris *et al*, 2008). Later, however, we identified sex dimorphism both in white matter morphometric abnormalities in ALC men and women (Ruiz *et al*, 2013), and also in the size of reward system regional volumes (Sawyer *et al*, 2017). In the present study, we sought to confirm our prior findings of lower FA in white matter in abstinent ALC men in an independent sample, and additionally, to compare these white matter abnormalities to those in abstinent ALC women. Based on our prior reports of sex dimorphism in alcoholic brains, we hypothesized that sex-related differences in white matter fiber tracts would vary between the ALC and NC groups.

## Materials and Methods

### Participants

Participants were solicited through online advertisements, newspaper advertisements, and flyers posted at participating institutions and in public places, e.g., churches, stores, etc. The study included 49 ALC participants (26 women), and 41 NC participants (22 women). They were right-handed, fluent English speaking individuals who ranged in age from 23 to 76 years. The groups were age balanced and had comparable socioeconomic backgrounds, and their ethnic distributions were similar (ALC: 1 American Indian, 7 Black, 41 White; NC: 1 Asian, 11 Black, 1 Hispanic, 28 White). The ALC participants met DSM-IV criteria for alcohol abuse or dependence, and had a minimum duration of five years of heavy drinking (21 or more drinks per week). As detailed in the Results section, all four subgroups performed comparably on neurobehavioral screening tests; the source data and code are provided in the Supplemental Dataset S1 and Supplemental Code S2. The research was approved by the Institutional Review Boards of Boston University School of Medicine, VA Boston Healthcare System, and Massachusetts General Hospital. Participants were reimbursed for their time and travel expenses.

### Inclusion and Exclusion Criteria

Selection procedures included structured interviews to determine education level, health history, and history of alcohol and drug use. Individuals were excluded from further participation if any source indicated one of the following: Korsakoff’s syndrome; HIV; cirrhosis; major head injury with loss of consciousness >30 minutes; stroke; epilepsy or seizures unrelated to alcoholism; Hamilton Rating Scale for Depression (HRSD; (Hamilton, 1960)) score over 18; schizophrenia; or electroconvulsive therapy. Additionally, we excluded individuals who failed screening for MRI safety (e.g., metal implants, obesity, pregnancy). Further exclusion of participants took place during the preprocessing of the MRI data, if they did not complete the MRI scan; showed excessive head motion and instability during the scan; or had a brain abnormality discovered by the scan.

The ALC individuals were included if they had a history of five or more years of heavy drinking (21 or more drinks per week), met Diagnostic and Statistical Manual (DSM-IV) (APA, 2000) criteria for alcohol abuse or dependence, and had been abstinent for at least four weeks before testing and scanning, with the exception of one man who drank the prior weekend, one man who drank three weeks before the scan, and one woman who drank six days before the scan. All three participants reported that the drinking occurred once, and they remained abstinent thereafter. A comprehensive psychiatric interview (described below) indicated that none of the participants had current drug abuse or dependence for any illicit substance including marijuana. Moreover, none of the participants had used illicit drugs for the four years preceding enrollment, with the exception of one ALC woman who stopped occasional marijuana use six months prior to testing, and one ALC man who stopped occasional marijuana use two months prior to testing.

### Evaluation of Drinking History

Drinking history was evaluated using a standardized alcohol use questionnaire (Cahalan *et al*, 1969) that yielded three measures: (1) length of sobriety (LOS); (2) duration of heavy drinking (DHD), i.e., the number of years consuming ≥21 drinks per week (one drink: 355 mL beer, 148 mL wine, or 44 mL hard liquor); and (3) a quantity frequency index, roughly corresponding to number of daily drinks (DD; one ounce of ethanol per drink). The DD measure evaluated the amount, type, and frequency of alcohol usage over the last six months (for NC participants), or over the six months preceding cessation of drinking (for ALC participants). To ensure that DD was comparable between ALC participants with complete abstinence, and four ALC who had quit but then drank occasionally (DD<3.0), we estimated DD based on the last six months of *heavy* drinking in the latter group. Thus, the measure consistently reflected DD across all ALC participants.

### Neurobehavioral and Psychiatric Evaluations

In order to minimize confounding effects from illicit drug use, psychoactive prescription drug use, and psychiatric comorbidity, participants were given a battery of screening tests, one of which was a computer-assisted, shortened version of the Diagnostic Interview Schedule (Robins *et al*, 2000) that provided lifetime psychiatric diagnoses according to DSM-IV criteria (APA, 2000). Additionally, tests of intelligence, memory, and affect were administered, including the Wechsler Adult Intelligence Scale, Third Edition (WAIS-III) for Full-Scale IQ (FSIQ), Verbal IQ (VIQ), and Performance IQ (PIQ); the Wechsler Memory Scale, Third Edition (WMS-III) for Immediate Memory, Delayed Memory, and Working Memory (Wechsler, 1997); and the HRSD to assess depression. Neurobehavioral and psychiatric evaluations were extensive and typically required six to nine hours over three or more days.

For neurobehavioral, psychiatric, and drinking variables, differences in diagnostic Group (ALC vs. NC) and Sex (men vs. women) were assessed using two-tailed Welch two sample t-tests on each of the measured indices in each category.

### Diffusion Imaging

Diffusion-weighted dMRI images were acquired on a 3 Tesla Siemens Tim Trio scanner to compute quantitative white matter diffusivity measures. A 12 minute 60-direction dMRI scan was obtained with 2x2x2 mm resolution at b=700 s/mm^2^ (T2 b0 images=10, TE=94 ms, TR=9800 ms, interleaved axial slices=64, FOV=256 mm, matrix=128x128, skip=0 mm, bandwidth=1860 Hz/pixel). For individual single-subject level and group-level comparisons of the dMRI processing stream, FSL v5.0 (Smith *et al*, 2004) Tract-Based Spatial Statistics (TBSS v1.1, www.fmrib.ox.ac.uk/fsl/TBSS) were used. We applied Threshold Free Cluster Enhancement (TFCE), a stringent control for Type I family-wise error rate that corrects for multiple comparisons and eliminates the need for setting an arbitrary cluster-forming threshold by merging voxel-wise statistics with local spatial neighbourhood information (Smith and Nichols, 2009). Details on the diffusion imaging processing stream are provided in Supplementary Methods.

We examined three contrasts: Interactions between Group and Sex, Group effects for men, and Group effects for women. Next, individual subjects’ average FA values were extracted from each cluster with significant Group by Sex interactions, and subsequent *ad hoc* analyses were performed, as described in the Supplementary Methods. In addition, we examined the relationships of FA from these clusters to drinking history and memory scores.

## Results

### Neurobehavioral, Psychiatric, and Drinking Measures

There were no significant differences between the ALC and NC participants with respect to age, education, WAIS-III, and WMS-III indices. There also were no significant Group by Sex interactions or main effects with respect to age, IQ, memory scores, depression, and education (Table 1). Although the depression scores for the ALC group were significantly higher than for the NC group (*p*<0.001), both groups’ scores were low (mean 3.9 vs. 1.0): HRSD scores of 8, 16, and 25 or above indicate mild, moderate, or severe depression, respectively (Zimmerman *et al*, 2013). The source data and code for these data are provided in the Supplemental Dataset S1 and Supplemental Code S2.

**Table 1.** Participant Demographic Characteristics and Drinking Histories.

|  | **Alcoholic Men** |  | **Alcoholic Women** |  | **Nonalcoholic Men** |  | **Nonalcoholic Women** |  |
| --- | --- | --- | --- | --- | --- | --- | --- | --- |
|  | **N = 23** |  | **N = 26** |  | **N = 19** |  | **N = 22** |  |
|  | Mean | (SD) | Mean | (SD) | Mean | (SD) | Mean | (SD) |
| Age (yrs) | 54.03 | (11.39) | 51.63 | (11.97) | 49.85 | (13.36) | 56.70 | (14.03) |
| Education (yrs) | 14.39 | (2.55) | 15.37 | (2.25) | 15.11 | (1.85) | 15.64 | (2.52) |
| FSIQ | 108.68 | (17.77) | 112.72 | (15.64) | 112.53 | (13.04) | 112.68 | (14.55) |
| VIQ | 110.64 | (16.49) | 114.76 | (14.71) | 113.79 | (13.18) | 112.64 | (16.14) |
| PIQ | 104.05 | (18.81) | 107.76 | (16.13) | 109.21 | (15.55) | 110.41 | (13.31) |
| IMI | 102.48 | (14.39) | 116.88 | (19.15) | 111.74 | (16.20) | 120.18 | (12.38) |
| DMI | 107.14 | (12.48) | 118.80 | (19.45) | 112.00 | (17.07) | 119.36 | (13.79) |
| WMI | 102.86 | (11.87) | 105.96 | (15.34) | 106.00 | (13.65) | 109.73 | (13.62) |
| HRSD** | 4.00 | (5.24) | 3.72 | (3.55) | 0.84 | (1.07) | 1.23 | (2.16) |
| DHD (years)* | 18.65 | (9.10) | 12.15 | (5.72) | 0.18 | (0.80) | 0.09 | (0.43) |
| DD (ounces/day)* | 15.64 | (11.95) | 8.16 | (7.34) | 0.42 | (0.69) | 0.19 | (0.34) |
| LOS (years) | 4.99 | (7.64) | 8.65 | (10.06) | NA | NA | NA | NA |

Abbreviations: Wechsler Adult Intelligence Scale measures are FSIQ: Full-Scale IQ; VIQ: Verbal IQ; PIQ: Performance IQ. Wechsler Memory Scale measures are: IMI: Immediate Memory Index; DMI: Delayed Memory Index; WMI: Working Memory Index. HRSD: Hamilton Rating Scale for Depression. WAIS-III and WMS-III scores were not available from one alcoholic woman and one alcoholic man, and WMS-III scores were not available from an additional alcoholic man. DHD: Duration of Heavy Drinking (years); DD: Daily Drinks (ounces of ethanol per day); LOS: Length of Sobriety (years). On average, alcoholic men drank 6.5 years longer ( $p<0.01$ ) and had 7.5 more daily drinks than women ( $p<0.02$ ). The difference between men and women in LOS was not significant ( $p=0.15$ ).
\*ALC men>ALC women, $p<0.05$ . \*\*ALC>NC, $p<0.001$ .

In order to account for any potential biases or confounding induced by sex differences in drinking histories, we performed a subanalysis (N = 81) for which ALC men and women had comparable drinking history (i.e., for which there were no significant differences). The TFCE corrected FA cluster volumes obtained from the interaction effect for this subgroup analysis overlapped with 98% of the cluster volumes obtained from the entire group, indicating the subgroup results were consistent with the entire group (as presented below).

### Fractional Anisotropy

As mentioned earlier, there were no significant results from the axial or radial diffusivity analyses. For FA however, three significant white matter tract clusters displayed Group by Sex interactions (*p*<0.05 using TFCE). The clusters were overlaid with a white matter atlas (Makris *et al*, 1999) and examined by a skilled neuroanatomist (N.M.). The anatomical locations are shown in Figure 1 and described in Table 2 (Figure S1 shows the non-thresholded contrasts.) The clusters included the genu and anterior parts of the body of the corpus callosum (CC2, CC3, & CC4), the left arcuate fasciculus and extreme capsule (AF & EmC) (Makris and Pandya, 2009), and the left superior longitudinal fasciculi (SLFII & SLFIII) (Makris *et al*, 2005). Of note regarding the corpus callosum, TFCE correction at first revealed two clusters (CC2 & CC3; 517 mm^3^) and (CC4; 4 mm^3^) separated by 6.1 mm. Our neuroanatomist (N.M.) determined that CC2, CC3, and CC4 were within the corpus callosum, and if we were to have used an adjusted threshold of *p*<0.06, the two clusters would have joined to form a single cluster. Therefore, the two clusters were combined into one 521 mm^3^ cluster.

**Figure 1.**
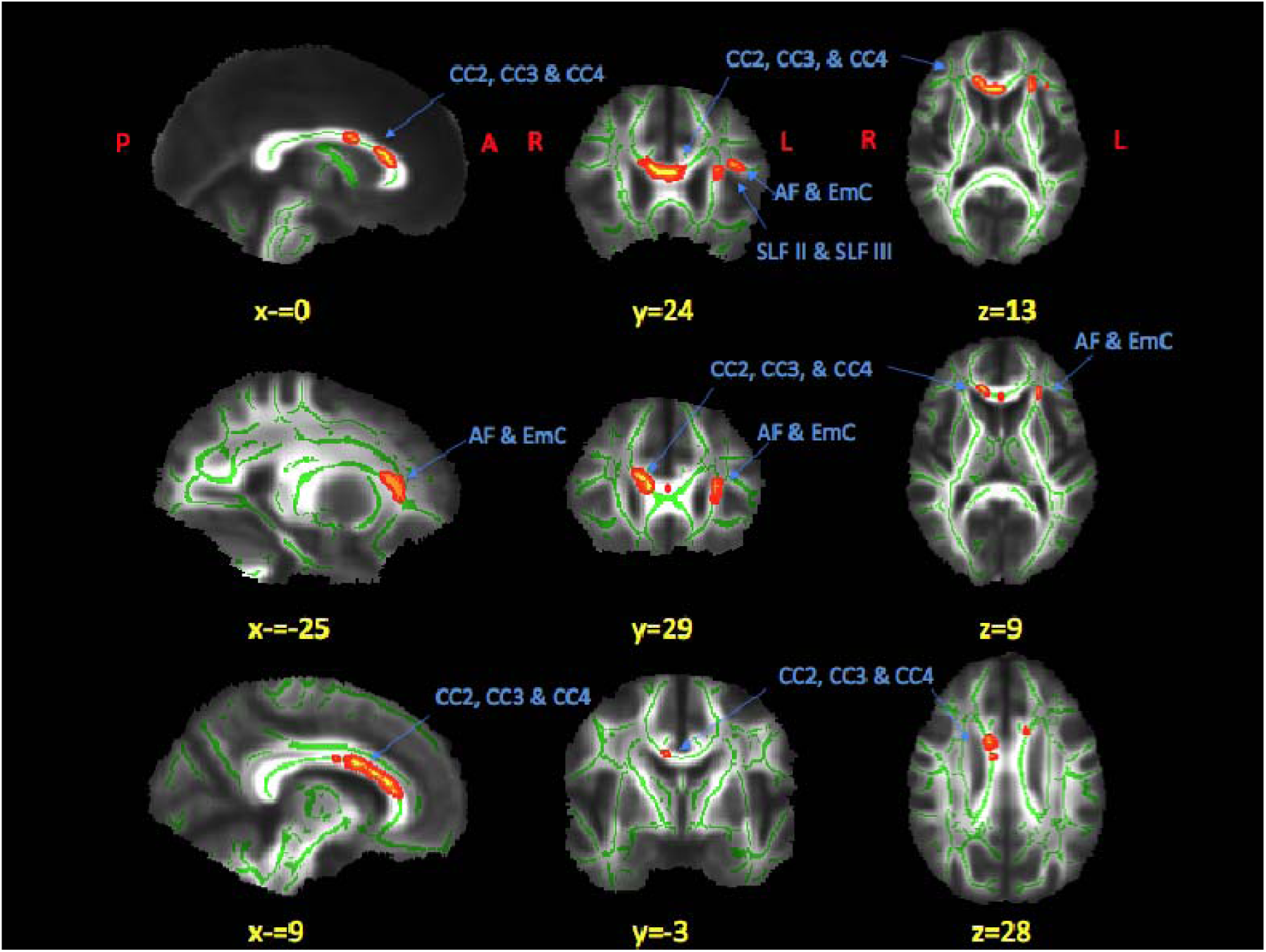
Significant Group by Sex interaction in FA differences in white matter. The maps represent significant (*p*<0.05, TFCE corrected) clusters from voxel-based FA Group by Sex interaction (red-yellow) in sagittal, coronal, and axial views. The green represents the skeletonized mean FA map of all of the subjects, which has been overlaid on a mean FA map of all the subjects. The red-yellow clusters represent regions with significant Group by Sex interactions. Three clusters were located as follows: the anterior portions and body of the corpus callosum (CC2, CC3, & CC4); the left superior longitudinal fasciculi (SLF II & SLF III); and the left arcuate fasciculus and extreme capsule (AF & EmC); see Figure 2 and Table S1 for FA levels.

**Table 2.** Significant Clusters from FA Comparisons. After threshold-free cluster enhancement multiple-comparison correction comparing Group by Sex interactions for FA differences, three clusters were identified as significant (Figure 1 for anatomical locations, Figure 2 for FA levels, and Table S1 for mean cluster values). Abbreviations: AF: arcuate fasciculus; SLFII & SLFIII: left superior longitudinal fasciculi; EmC: extreme capsule; CC2 and CC3: anterior portions of the corpus callosum; CC4: body of the corpus callosum; PTR: posterior thalamic radiation. TalX, TalY, and TalZ represent the Talairach coordinates of the most significant voxel in the cluster. The coordinates were translated to the Talairach coordinates using Yale’s BioImage Suite (Papademetris *et al*, 2006).

| Contrast | Cluster # | Cluster size (mm<sup>3</sup>) | TalX | TalY | TalZ | Annotation | Hemisphere |
| --- | --- | --- | --- | --- | --- | --- | --- |
| Group by Sex interaction | 1 | 40 | -33 | 22 | 17 | SLF II & SLF III | Left |
| Group by Sex interaction | 2 | 69 | -24 | 26 | 10 | AF & EmC | Left |
| Group by Sex interaction | 3 | 521 | 0 | 22 | 13 | CC2, CC3 & CC4 | Bilateral |
| FALC vs FNC | ns | ns | ns | ns | ns | ns | ns |
| MALC vs MNC | 4 | 271 | -29 | -56 | 15 | PTR | Left |

The relationships between groups with mean FA in these clusters are shown in Figure 2 and Table S1. Similar interactions were observed for all three clusters: ALC men had lower FA than NC men, while ALC women had higher FA than NC women (all Welch’s t-test *p*<0.05).

**Figure 2.**
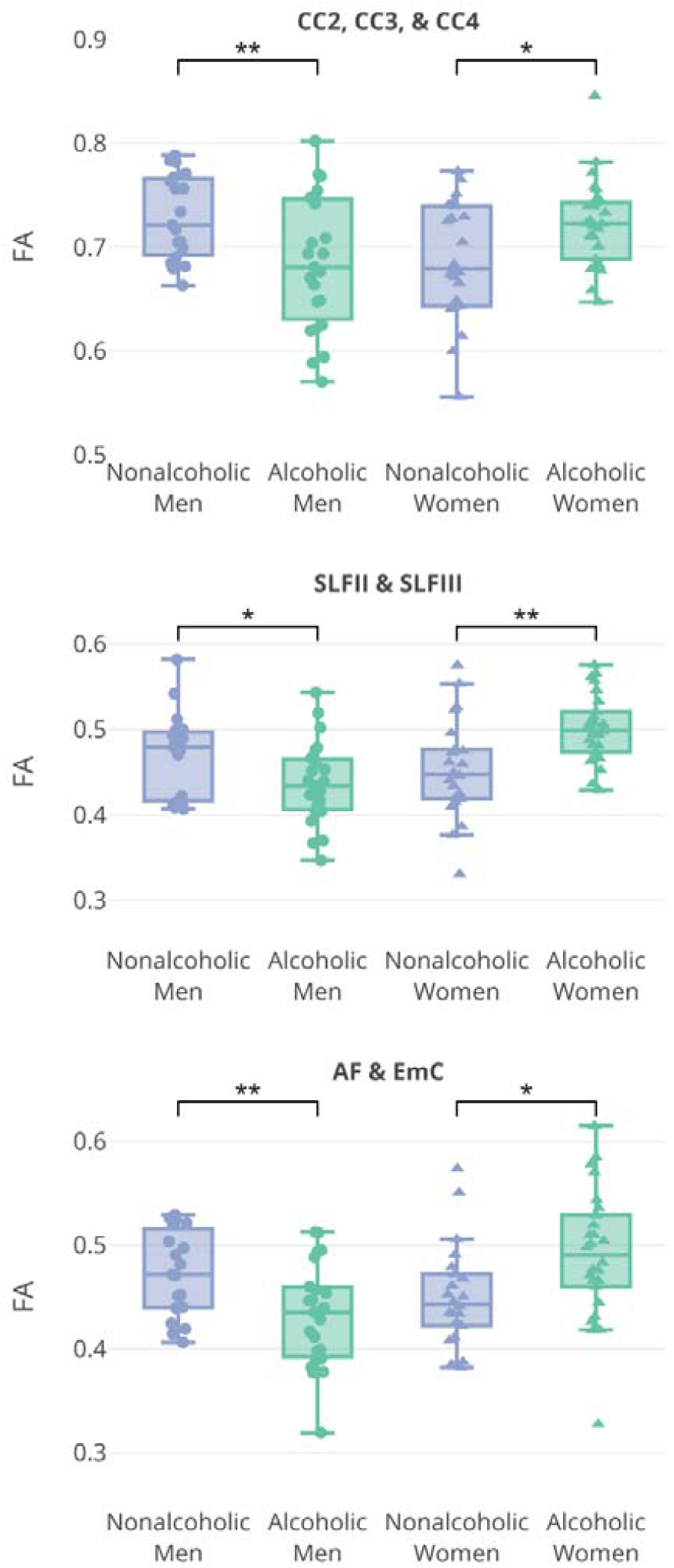
The relationship of alcoholism and fractional anisotropy (FA) differs for men and women. In a significant two-way interaction of Group by Sex, three clusters were identified after threshold-free cluster enhancement correction for multiple comparisons (*p*<0.05). ALC men had lower FA than NC men, while ALC women had higher FA than NC women (\**p*<0.05; ** *p*<0.01). The three clusters were located as follows: the anterior portions and body of the corpus callosum (CC2, CC3, & CC4); the left superior longitudinal fasciculi (SLF II & SLF III); and the left arcuate fasciculus plus extreme capsule (AF & EmC); see Figure 1 for cluster locations and Table S1 for mean values.

As a secondary analysis, in addition to assessing Group by Sex interaction effects, we included an analysis of the group differences for men and women separately in our TBSS model (Table 2). For women, no clusters were significant after TFCE correction. For men, one cluster was identified, as shown in Figure S2 and Figure S3. This cluster was superior to the posterior horn of the left lateral ventricle, and included the posterior thalamic radiation (PTR). For the PTR cluster, alcoholic men had 0.07 lower FA than nonalcoholic men (Table S1).

### Behavioral Correlations with Regional FA Measures

We examined the relationship of our three drinking history measures (DHD, DD, and LOS) to FA for the three interaction clusters we identified (Table S2X-S2X). Further, we determined how these relationships differed for men and women by examining the interaction of Sex with each drinking history measure (Table S2X-S2X). A significant interaction with Sex was found for the CC2, CC3 & CC4 cluster: FA was positively associated with LOS for men, while no significant relationship was found for women (Figure 3 and Table S2C).

**Figure 3.**
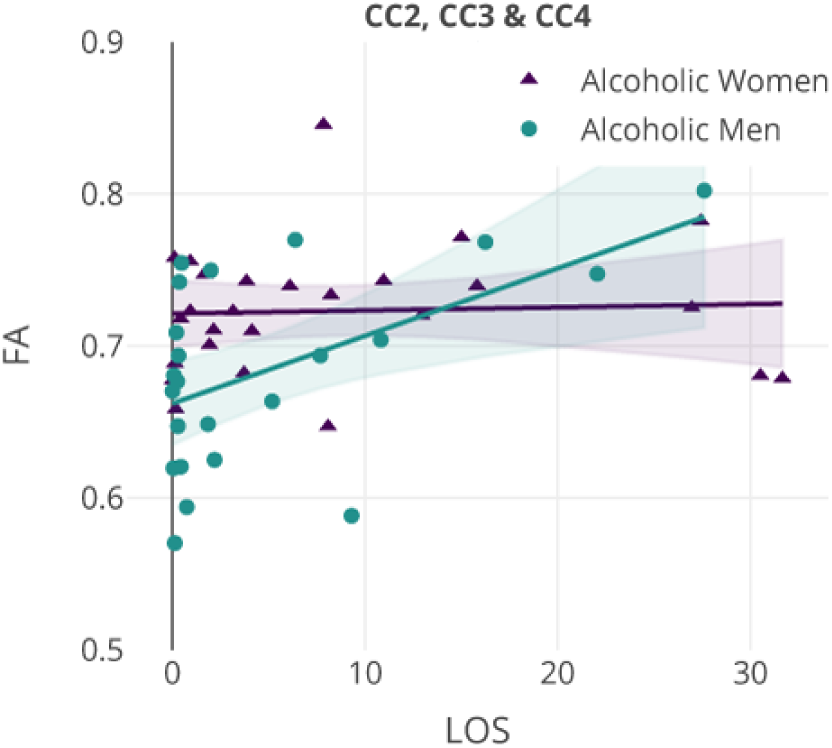
Fractional anisotropy is significantly related to drinking history differently in men and women with alcoholism.The CC2, CC3 & CC4 cluster showed a significant interaction effect in the relationship between Length of Sobriety (LOS) and Sex; see Table S2. Regression lines and 95% confidence intervals are displayed separately for men and women. Women are displayed with triangles and men are displayed with circles.

In addition, for the total sample of 90 participants, we examined the relationships of the mean FA of each cluster (Table S3) to (1) age at scan, and (2) memory scores (IMI, DMI, and WMI). FA declined with age by 0.001 per year in the SLF II & SLF III clusters (*p*<0.05), and 0. 002 per year, both in the AF & EmC cluster, and in the CC2, CC3 & CC4 cluster (*p*<0.001). IMI was associated with higher FA for the CC2, CC3, & CC4 cluster (*p*<0.05), while DMI and WMI were not significantly related to FA in any cluster. A significant interaction of Group and Sex was observed with Age for the CC2, CC3, & CC4 cluster (*p*<0.05), where ALC men (*p*<0.05), NC men (*p*<0.01), and NC women (*p*<0.001) had negative relationships with Age, while ALC women did not (Table S3). No significant Group or Sex interactions were observed for the relationship of FA for any of the three clusters with IMI, DMI, or WMI.

## Discussion

While the effects of long-term chronic alcoholism on the brain have been examined in neuroimaging studies, most have included only or mainly men, or did not differentiate between men and women in a mixed sample. These studies generally have reported brain atrophy, cortical thinning, and smaller volumes of the frontal lobes, hippocampus, thalamus, pons, and cerebellum, with increases in the ventricles and cerebrospinal fluid in the men. DTI abnormalities reported in those prior studies have mainly encompassed fronto-cerebellar circuits, genu and splenium of the corpus callosum, the centrum semiovale, the superior longitudinal fasciculus, the external capsule, the fornix, and the cingulum (Harris *et al*, 2008; Pfefferbaum *et al*, 2007; Pfefferbaum and Sullivan, 2002,2005).

We had previously examined differences in cerebral white matter integrity between abstinent ALC and NC men (Harris *et al*, 2008). In the present study, we aimed to examine sex-specific differences in the impact of alcoholism, and potentially underlying sex dimorphic brain differences in the predisposition to alcoholism. The results showed sex-related abnormalities in the white matter tracts of the alcoholic individuals. The abstinent alcoholic men had white matter FA deficits consistent with our prior study, but the abstinent ALC women did not. These distinct patterns of white matter anomalies suggest a different underlying neural basis for sex-specific propensity (i.e., differential risk factors) and/or sequelae to long-term alcoholism, and have implications for further investigation of possible sex-specific approaches to prevention and treatment.

While the exact nature of pathophysiological white matter changes underlying the observed differences in the current study can not be inferred from DTI results alone, recent findings using nonhuman animal models of alcoholism may help to explain alcohol induced white matter deficits. Animal studies have provided morphological evidence that chronic alcohol exposure can cause damage not only to nerve cells but also to nerve fibers (Luo *et al*, 2014, 2017). Further confirming a link between morphological changes and neuronal damage, Luo and colleagues (Luo *et al*, 2017) have shown that nerve fiber changes in male rats with chronic alcohol exposure are detectable by DTI concurrent with changes in the level of N-Acetylaspartic acid (NAA), a sensitive surrogate for neuronal damage (Moffett *et al*, 2007). Reduction of NAA in the brains of alcoholics has been reported in humans (Jagannathan *et al*, 1996; Meyerhoff *et al*, 2004), and moreover, NAA loss may be recoverable and can increase with continued abstinence (Parks *et al*, 2002). Furthermore, in another study using electron microscopy, Luo and colleagues had shown patterns of axonal shrinkage and demyelination in the brainstem of male rats with chronic alcohol exposure (Luo *et al*, 2014).

In the present study, sex differences in the relationship of FA with alcoholism were observed for three clusters: the anterior and middle portions of the corpus callosum; the arcuate fasciculus and extreme capsule; and the superior longitudinal fasciculi II and III (Figure 1). As shown in Figure 2, ALC men had lower FA than NC men, while ALC women had higher FA than NC women. Parallel gender dimorphism was reported in left frontal cortical thickness in adolescent binge drinkers (Squeglia *et al*, 2012), suggesting that these gender differences may pre-date long-term alcoholism effects, and may reflect differential pre-existing factors, such as personality, prodromal psychiatric comorbidity, or motivational factors for drinking. In other words, it may be that drinking problems arise in men and women for different reasons (Nicolai *et al*, 2012; Ruiz and Oscar-Berman, 2015; Squeglia *et al*, 2012), and those associated risk factors could be manifested by differential prodromal FA abnormalities. The dominant motivation for men to drink may be to enhance sensation and social function, whereas drinking to cope with negative emotions may have a more nuanced relationship with sex (Kuntsche *et al*, 2006). Similarly, the motivation to drink may be differentially associated with personality factors (Mosher Ruiz et al, 2017). Additionally, there is a high prevalence for psychiatric conditions in conjunction with alcoholism. While we excluded participants who met DSM-IV criteria for depression, stressful life events could differentially contribute to the onset of heavy drinking (Keyes *et al*, 2011; Nolen-Hoeksema and Hilt, 2006). The prevalence of these conditions is higher in women than men (Khan *et al*, 2013). In contrast, another psychiatric comorbidity, antisocial personality, has been observed in conjunction with alcoholism more often in men (Compton *et al*, 2005; Oscar-Berman *et al*, 2009). Interactions of sex with alcoholism-related consequences and risk factors may be associated with unique brain regions. Although we observed the same direction of effects for all three of the pertinent DTI clusters, each could have different psychosocial impacts.

For the first cluster, the anterior corpus callosum conveys interhemispheric signals across prefrontal brain regions (Makris *et al*, 1999). In a previous study (Sawyer *et al*, 2017), the volume of dorsolateral prefrontal cortex was reported to be smaller in ALC men than NC men, while the opposite was observed for women, perhaps reflecting sex differences in FA effects. Further, the volume of the mid-anterior corpus callosum was smaller in ALC men than NC men (Ruiz *et al*, 2013). Considering the inherently specialized functions of the two human frontal lobes, the axonal abnormalities we observed in the corpus callosum could be tied to alcoholism-related abnormalities observed in personality (Westlye *et al*, 2011) and psychiatric conditions (White *et al*, 2008).

For the second cluster, a significant alcoholism Group by Sex interaction was observed in the anterior portion of the arcuate fasciculus and extreme capsule. While the arcuate fasciculus is thought to conduct signals primarily between Broca’s area and Wernicke's area, this interpretation remains unsettled (Rilling *et al*, 2008), and recent research suggests broader connectivity (Dick and Tremblay, 2012). While one might predict that the reduced connectivity observed in ALC men could impart a reduction in language ability, we did not observe one, nor have language deficits been widely reported (Oscar-Berman *et al*, 2014). The cluster also might include the extreme capsule, which may be involved in long-term semantic memory. However, caution is advised when interpreting dMRI results for this tract (Dick and Tremblay, 2012).

For the third cluster, we observed a similar pattern of FA abnormalities for a region including portions of two major longitudinal pathways, SLF II and SLF III, which have been implicated in diverse functions including working memory (Dick and Tremblay, 2012). Working memory deficits in alcoholism have been well established (Oscar-Berman *et al*, 2014), but further research could clarify the relationship of these abnormalities to sex and to pre-existing risk factors.

By examining the amount and duration of drinking, along with the duration of abstinence, in relation to the present dMRI findings, we can begin to suggest some consequences. We found that LOS was correlated with higher FA of the callosal white matter in men, but not for women (Figure 3). This interaction could indicate more recovery for men, considering our observation that the men with short sobriety periods had lower FA than women with short sobriety periods, but the opposite was true for long sobriety periods. Interestingly, in a recent study (Monnig *et al*, 2015) on a large sample of non-abstinent, heavily drinking alcoholics (324 participants, 30% female), more frequent drinking (a close match to our estimates of DD during the period of heavy drinking) was reported to contribute to lower FA in ALC women but not ALC men – even though both sexes were similar in terms of demographics, history of drinking, and drinking severity. Previous research found lower FA values in ALC women as compared to ALC men, when both groups were matched by the amount of lifetime alcohol consumption and length of abstinence (Pfefferbaum *et al*, 2009). The ALC men and women in our study were not matched for either of these variables; however, our findings were consistent when we analyzed a subgroup that included men and women with comparable drinking histories.

As aging also has been associated with a decline in the integrity of white matter microstructure (Salat *et al*, 2005; Yang *et al*, 2016), we examined the relationship of age to the three clusters with significant Group by Sex interactions. Age was correlated with lower FA in each cluster, at levels comparable to those previously reported (-0.002 to -0.001 per year).

## Limitations

While we found strong evidence for alcoholism sex differences in white matter integrity, this study had several limitations. First, as noted above, our sample of ALC women had shorter (by 6.5 years) and less severe (7.5 fewer DD) drinking histories than the ALC men. However, when analyzed subgroups with comparable drinking histories, we found that the clusters remained significant. Second, we did not counterbalance for tobacco usage, which may interact with alcoholism (Durazzo *et al*, 2013; Luhar *et al*, 2013). Nor did we counterbalance recruitment by the participants’ family history of alcoholism, so the differences in FA we observed may be related to heritable traits. Finally, the cross-sectional nature of this study hinders causal interpretation; that is, we cannot determine whether the FA differences predated the onset of drinking, or were caused by heavy drinking and associated sequelae. In any case, the findings presented here extend the results found in ALC men to a sample of women alcoholics, and broaden the knowledge base of sex differences in relation to the psychobiology of alcohol use disorders.

## Conclusions

This DTI study provides evidence in support of a growing understanding of the nature of sex dimorphism in brain and behavioral abnormalities associated with alcoholism. Specifically, we found differences between alcoholic men and women in white matter integrity, evidenced by lower FA values in ALC men and higher FA in ALC women. The white matter abnormalities in men observed in our study support previous literature on alcoholism, specifically those studies indicating damage to cortical and limbic brain regions that comprise a system essential for normal emotional functioning, and modulating the effectiveness of positive and negative reinforcement in human behavior. Future research is needed to explore whether sexual dimorphism in white matter microstructure may implicate underlying differences in the neurobehavioral liabilities for developing alcohol abuse disorders, or for sequelae following abuse.

## Acknowledgements

We thank Elinor Artsy, Sheeva Azma, Julie Howard, Diane Merritt, Alan Poey, Daniel Salz, Yulia Spantchak, Trinity Urban, Susan Mosher Ruiz, Maria Valmas, and Robert Zondervan for assistance with recruitment, assessment, or neuroimaging of the research participants.

## Supplemental Methods

For each subject, the dMRI scans were visually inspected in all 60-direction volumes. After skull-stripping with the Brain-Extraction Tool (BET), we used the FMRIB Diffusion Toolbox v2.0 (Behrens *et al*, 2003) to perform motion and eddy current distortion corrections (Jenkinson *et al*, 2002; Jenkinson and Smith, 2001; Smith, 2002), and to generate a diffusion tensor for each voxel, calculated using a least squares fit of the tensor model to the dMRI data. From the diffusion tensors, the eigenvalues of each tensor (which represent the magnitude of the three main diffusion directions) were calculated for each voxel. Maps of axial diffusivity, radial diffusivity, and FA were created for each participant. There were no statistically significant results from the axial or radial diffusivity analyses we specified. The FA maps served as the primary measure of white matter microstructural integrity.

In order to prepare the single subject level data for group level analyses, all of the maps were aligned to the standard 1×1×1 mm^3^ MNI152 brain, using nonlinear registration and subsequently skeletonized with a threshold of FA > 0.2 to reduce the likelihood of partial voluming with the bordering gray matter and ventricular cerebrospinal fluid. To co-register the core white matter pathways among all of the subjects in this analysis, the following procedure was employed: A nonlinear registration was performed in order to coregister or align all FA images from all subjects to a predefined FA template image; then the FSL-based FA template (i.e., the target image) was derived from an averaged dataset of 58 FA maps from all healthy subjects, both male and female. The FA template was also in the standard 1×1×1 mm^3^ MNI152 space. Using TBSS, the calculated nonlinear transformation was applied to the estimated pathways for each individual subject, so all subjects were coregistered to MNI152 space for group level analysis. These maps were then binarized. The thresholded, normalized and nonlinearly warped and binarized maps were then summed across subjects to produce a group average probability map.

Group-level differences in dMRI data were assessed by standard Tract-Based Spatial Statistics (TBSS v1.1, www.fmrib.ox.ac.uk/fsl/TBSS) analyses (Smith *et al*, 2006). To determine group-level differences in FA between the ALC and NC groups, whole brain voxel-wise statistical analyses were performed on the skeletonized data using FSL randomise (Winkler *et al*, 2014), a nonparametric permutation inference tool with threshold-free cluster enhancement (TFCE), which corrects for Type I error (Woo *et al*, 2014). The TFCE multiple comparison correction within TBSS analysis was performed on the comparison results by voxel-wise permutation (5000 permutations) testing, with a cluster-wise significance threshold of *p*<0.05.

Clusters with significant Group by Sex interactions were extracted, and further *ad hoc* analyses were performed using R 3.3.2 (Daróczi and Tsegelskyi, 2015; Dowle *et al*, 2015; R Core Team, 2016; Robinson, 2014; Sievert *et al*, n.d.; Xie, 2013). Two-tailed Welch two-sample t-tests were used to compare the cluster FA values of the ALC to NC groups for men and women separately. The FA values for each cluster also were examined for relationships with age, and for immediate, delayed, and working memory for all participants, and interactions of those measures with group and sex. For the ALC group, interactions of drinking history (DHD, DD, and LOS) with sex were examined, followed by the direct relationships of drinking history to FA where significant interactions were not present.

---

**Table S1.** Fractional anisotropy values for significant clusters. The mean and standard deviation (SD) fractional anisotropy values are listed separately by group and sex for each cluster. The first three clusters (CC2, CC3 & CC4: the anterior portions and body of the corpus callosum; AF & EmC: the left arcuate fasciculus and extreme capsule; and SLFII & SLFIII: the left superior longitudinal fasciculi) were obtained using a TBSS contrast examining the interaction of Group by Sex, with TFCE correction. The last cluster (PTR: posterior thalamic radiation), was obtained from a TBSS contrast comparing ALC men with NC men.

| Clusters | Alcoholic Men |  | Alcoholic Women |  | Nonalcoholic Men |  | Nonalcoholic Women |  |
| --- | --- | --- | --- | --- | --- | --- | --- | --- |
|  | Mean | (SD) | Mean | (SD) | Mean | (SD) | Mean | (SD) |
| SLFII & SLFIII | 0.44 | (0.05) | 0.50 | (0.04) | 0.47 | (0.05) | 0.45 | (0.06) |
| AF & EmC | 0.43 | (0.05) | 0.49 | (0.06) | 0.47 | (0.04) | 0.45 | (0.05) |
| CC2, CC3, & CC4 | 0.68 | (0.06) | 0.72 | (0.04) | 0.73 | (0.04) | 0.69 | (0.06) |
| PTR | 0.54 | (0.08) | 0.59 | (0.05) | 0.60 | (0.04) | 0.59 | (0.03) |

**Table S2.**
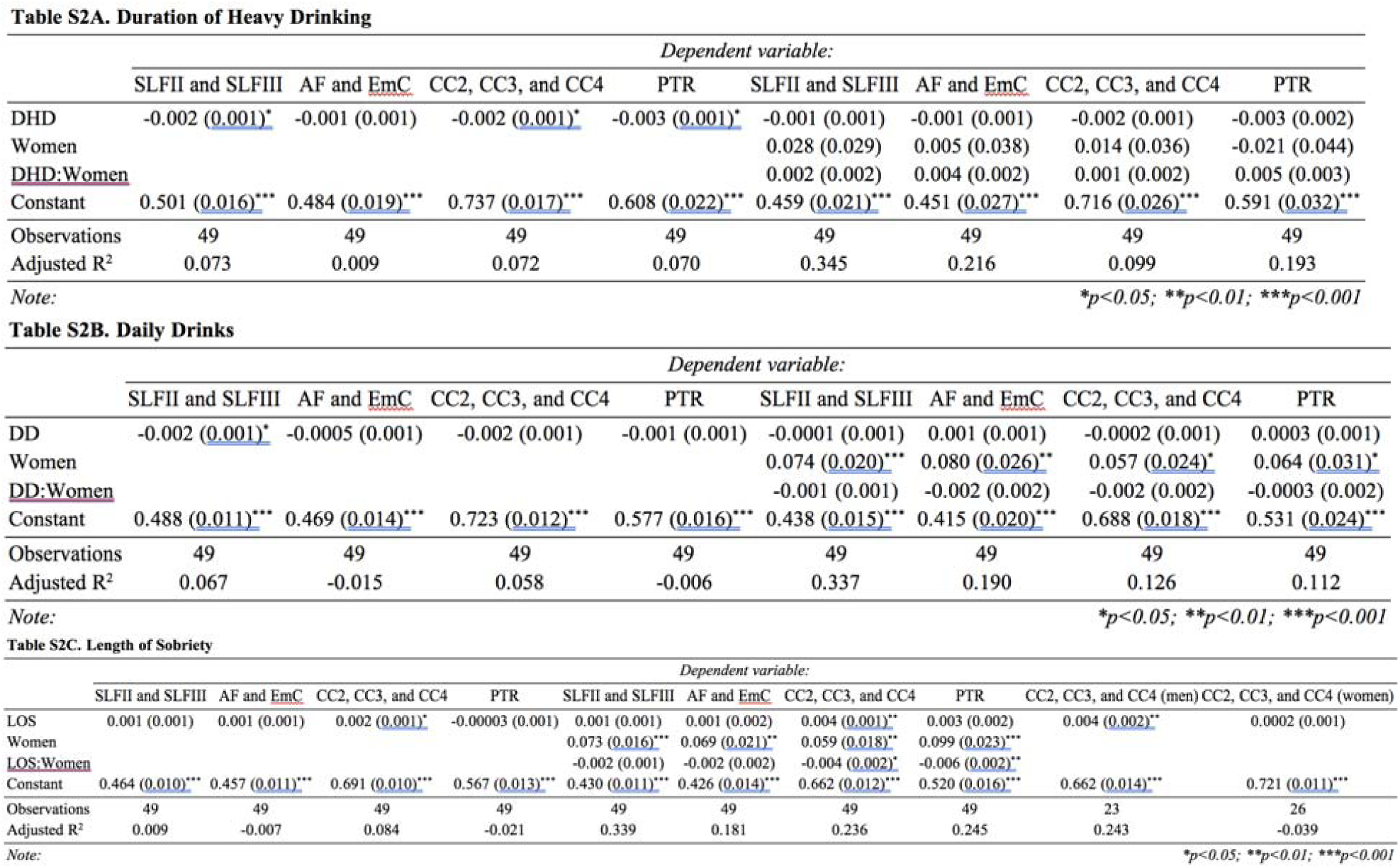
Regression reports for cluster FA predicted by drinking history measures. Linear regression results are reported for the relationship of fractional anisotropy to the three measures of drinking history: duration of heavy drinking (DHD; years), daily drinks (DD; ounces of ethanol per day), and length of sobriety (LOS; years): Tables S2A-S2C. Models examining interactions of these drinking measures with Sex are reported as well: Tables S2A-S2C. Table S2C shows the relationship of LOS with the CC2, CC3, & CC4 cluster FA for alcoholic men and women separately.

**Table S3.**
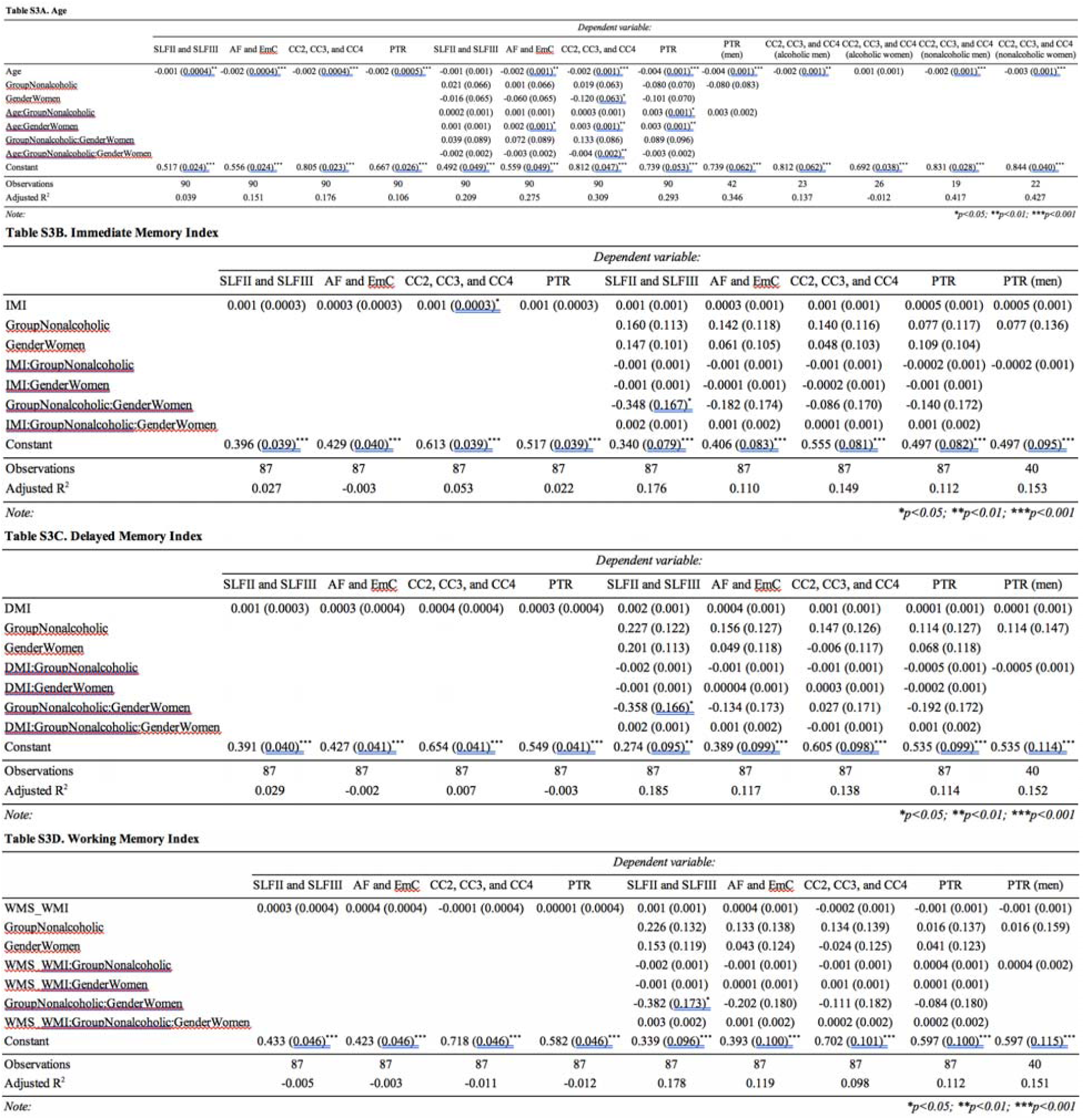
Regression reports for cluster FA predicted by age and memory scores. Linear regression results are reported for the relationship of fractional anisotropy to age and the three measures of memory from the Wechsler Memory Scale, Third Edition: Immediate Memory Index (IMI), Delayed Memory Index (DMI), and Working Memory Index (WMI): Tables S3A-S3D. Models examining interactions of these drinking measures with Sex and Group are reported as well.

**Figure S1.**
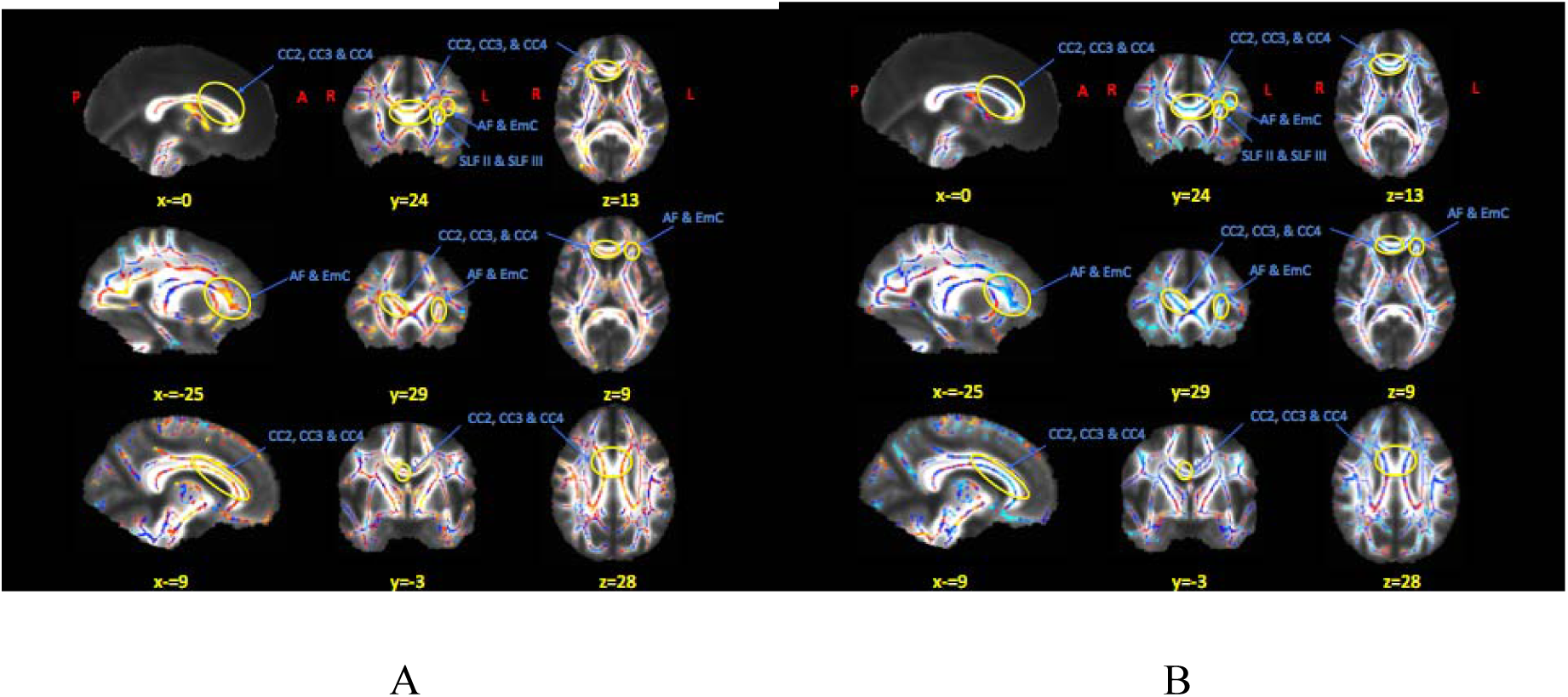
Group by Sex interaction in fractional anisotropy differences in white matter. The non-thresholded contrast of parameter estimate FA map is displayed for (A) alcoholic men vs. nonalcoholic men, and (B) alcoholic women vs. nonalcoholic women, displayed in in sagittal, coronal, and axial views. Blue-light blue represents regions where the parameter estimate for the FA values in corresponding ALC group is higher than the NC group. Red-Yellow colors represent regions where the parameter estimate for the FA values in corresponding ALC group is lower than the NC group. Yellow circles indicate regions where the Group by Sex interaction TFCE analysis revealed significant clusters (Figure 1). The three clusters were located as follows: the anterior portions and body of the corpus callosum (CC2, CC3, & CC4); the left superior longitudinal fasciculi (SLF II & SLF III); and the left arcuate fasciculus and extreme capsule (AF & EmC); see Figure 2 for individual FA levels.

**Figure S2.**
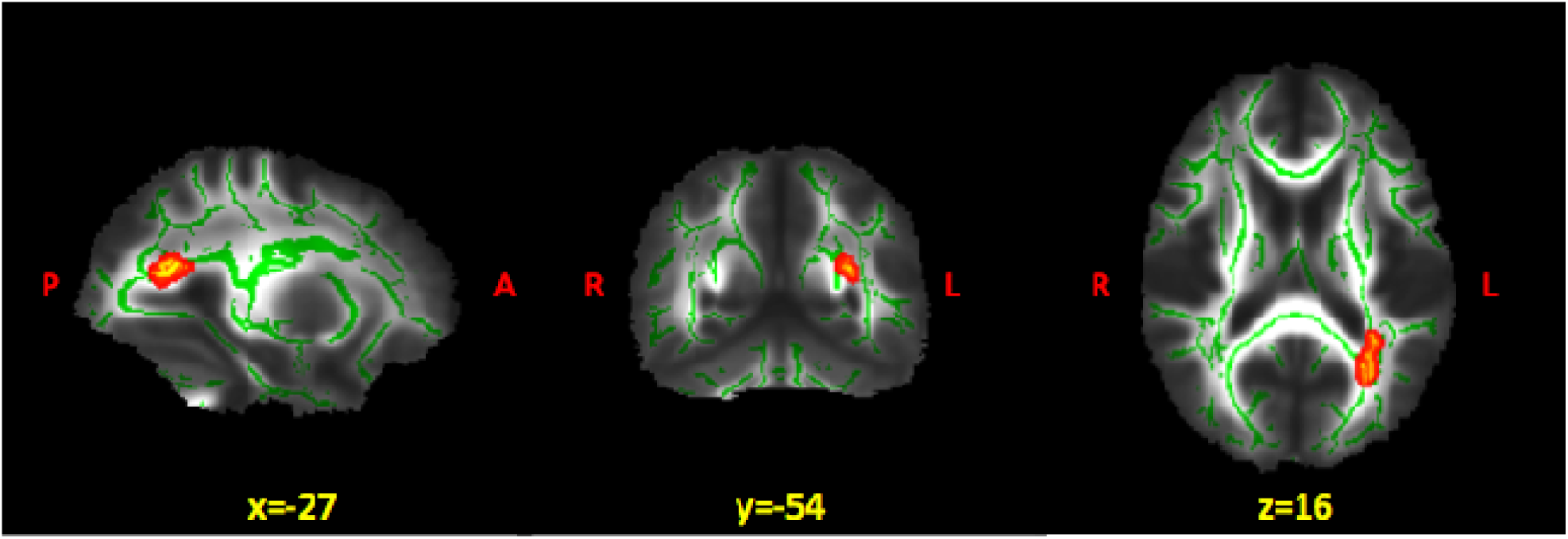
Alcoholic men had lower fractional anisotropy than nonalcoholic men in the PTR cluster. The maps represent significant (*p*<0.05, TFCE corrected) clusters from voxel-based FA value comparisons in alcoholic men vs. nonalcoholic men (in red-yellow) in sagittal, coronal, and axial views. The green represents the skeletonized mean FA map of all of the subjects, which has been overlaid on a mean FA map of all the subjects. The red-yellow clusters represent regions with significant effect. One cluster was identified, in the posterior thalamic radiation (PTR); see Figure S3 for individual FA levels.

**Figure S3.**
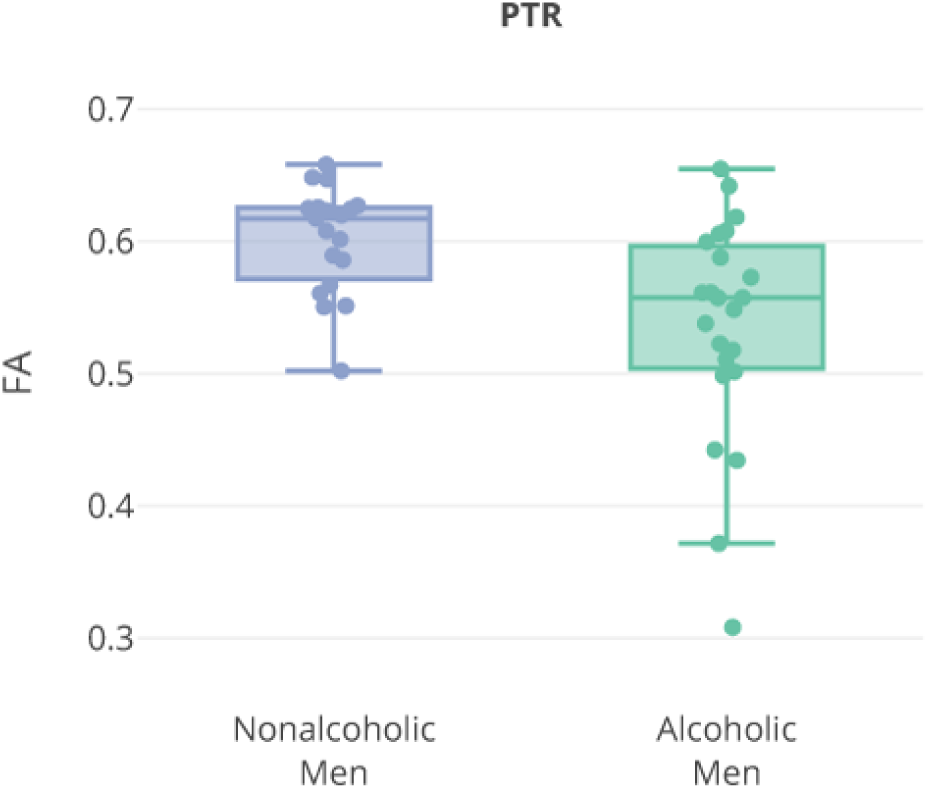
Alcoholic men had lower fractional anisotropy than nonalcoholic men in the PTR cluster. When examining the effect of alcoholism group for men and women separately, one clusters was identified after TFCE correction for multiple comparisons (*p*<0.05). For the posterior thalamic radiation (PTR), alcoholic men had 0.06 lower FA than nonalcoholic men; see Figure S2 for cluster locations.

